# Juvenile honest food solicitation and parental investment as a life history strategy: a kin demographic selection model

**DOI:** 10.1101/240416

**Authors:** József Garay, Villő Csiszár, Tamás F. Móri, András Szilágyi, Zoltán Varga, Szabolcs Számadó

## Abstract

Parent-offspring communication remains an unresolved challenge for biologist. The difficulty of the challenge comes from the fact that it is a multifaceted problem with connections to life-history evolution, parent-offspring conflict, kin selection and signalling. Previous efforts mainly focused on modelling resource allocation at the expense of the dynamic interaction during a reproductive season. Here we present a two-stage model of begging where the first stage models the interaction between nestlings and parents within a nest and the second stage models the life-history trade-offs. We show in an asexual population that honest begging results in decreased variance of collected food between siblings, which leads to mean number of surviving offspring. Thus, honest begging can be seen as a special bet-hedging against informational uncertainty, which not just decreases the variance of fitness but also increases the arithmetic mean.

**Author Summary:** Parent-offspring communication is a fascinating problem that captures the attention of scientist and layman alike. Parent-offspring interaction is the first interaction with non-self in the life of most young birds and mammals. The future life success of such young animals crucially depends on how successful they are in interacting and communicating with their parents. This communication has different functions: it is important for the offspring to solicit food from the parent and it is important for the parent to be informed about the state (hunger level) of the offspring. There is an optimization problem on top of this level: the parent has to ‘decide’ what part of the available resources should be allocated to the offspring and what part should she keep for herself. Here we show in a probabilistic model that the honest phenotype -where offspring beg only if they are hungry-has a greater growth rate than a selfish type - which begs regardless of its hunger level. This result holds in asexual populations; here honesty serves as a reduction uncertainty for the parents. The improved decision making of the parents -in turn-increases the survival of the offspring as well.

## 1. Introduction

Parent-offspring conflict stays in the spotlight of biological investigations [1,2,3,4]. Begging behaviour where offspring solicit food from their parents can be found in many species ranging from insects to birds (see reviews [5,6]). Solicitation appears to be honest in general, as much as needier offspring beg more intensively [6]. Such honest begging is perplexing at first. Trivers [7] in his seminal work predicted that there is a conflict of interest between parent and offspring regarding the ratio of resources allocated to the offspring. If there is indeed a conflict of interest between parent and offspring, then offspring are expected to solicit more food than the parental optimum would be. Costly signalling was proposed as a solution to this problem [8,9,10,11]. These models argue that offspring solicitation must be costly, and this cost should remove conflict of interest between parent and offspring by moving the offspring’s optimal resource allocation into the position of the parental optimum [9,10,11].

While costly signalling models proposed a potential solution to the problem, there is still an ongoing debate about the validity of the assumptions of the costly signalling models and about the potential information content of begging calls [12,13,14,15,16]. These debates are difficult to resolve for several reasons. One of them is that begging is a special type of a conflict situation: (i) it is within the family, hence interactions are non-random; (ii) genes playing against their own copies [17]. Accordingly, parent-offspring conflict potentially violates two basic assumptions of game theoretical models: (i) random interaction of players, (ii) the idea of independent players.

On top of these problems there are at least two kind of optimization problems: (i) the first one is a resource allocation problem between parents and offspring with an element of uncertainty about the quality of the offspring; (ii) and the second one is a dynamic temporal problem of food allocation between offspring with an element of uncertainty about the satiation of the offspring (hunger). Roughly speaking, these two optimization problems correspond to the strategic vs. tactical levels of the situation. On the strategic level the question is how to allocate resources between parent and offspring based on the quality of the offspring and the quality of environment in order to achieve the strategic goal of the parents (i.e. phenotype fitness maximalization). For example, in good years parents can raise all the offspring, but they still might want to allocate more food to high quality offspring; on the other hand, in bad years the parents might want to raise only high-quality offspring and sacrifice the low-quality ones. It is worth to note that this first level is a conceptual level and parents never have to make this decision during their lives (i.e. what part of the overall resources should go to the offspring). Second, parents have to make tactical decisions in order to achieve their strategic goals, these are the actual decisions parents will make during their lifetime. They have to decide which offspring to feed at any given feeding bout so that (i) they receive the overall amount food optimal for the parent, and that (ii) those offspring survive that serve the interest of the parent. Figure 1 shows the relation of these strategic and tactical levels and the corresponding trade-offs.

**Figure 1.**
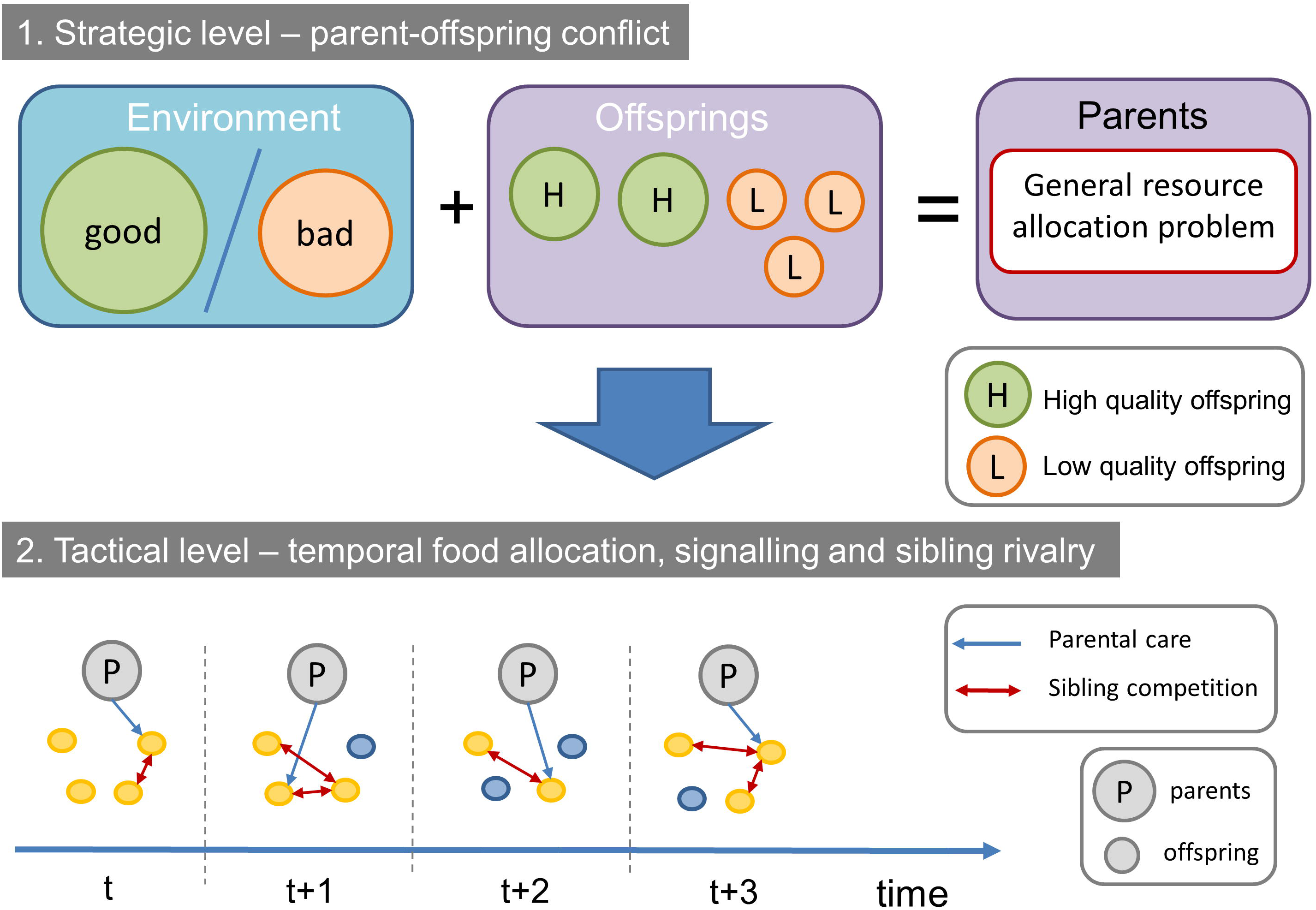
Strategic vs. tactical levels of the parent-offspring conflict during the reproductive period.

Note that these problems are obviously inter-related. The strategic goals cannot be achieved without good tactical decisions. Moreover, parents may not have all the information at the beginning of the reproductive season to set their strategic goals: they may not know the quality of the environment or the offspring or both. These strategic goals might even change during the season, the environment might be better than expected or some offspring might be eaten by predators, etc. However, we think that it is still important to differentiate between these problems on a conceptual and modelling level.

Since there are more than one optimization problems, parents are expected to gather information on all important aspects. We argue that the current debate in the literature is mostly due to the fact that these optimization problems are confused. Begging signals are complex, and it is highly likely that they contain information on both problems. How parents use this information is another, interesting question. Parents can use the information selectively, depending on the environment [1], ignoring one or the other component (which, of course corresponds to the different strategic goals: e.g. either raise all the offspring or raise only the best).

Here we present a minimal model to investigate these optimization problems. This model is a two-stage process where the first model investigates the tactical decisions of the parents in the temporal dimension, whereas the second model investigates the strategic resource allocation problem in the light of the results of the first model. Our main goal is to investigate the effects of honest signalling on the tactical level and to investigate how it interacts with the strategic goals of the parents.

Here we investigate an asexual population in order to keep our model analytically tractable. This means that the phenotype includes both the adult and the juvenile stage, hence these stages are not different players. In other words, we investigate a two-dimensional life history strategy (e.g. [17]) that includes both what an individual does at its juvenile and adult stage. The different players in our model are the different phenotypes: honest vs. dishonest. However, these phenotypes do not play against each other during the reproductive stage due to the structure of the game (i.e. interactions are within the family) and the nature of reproduction (i.e. like beget like). While honest and dishonest types do not compete in the family, yet they compete on the population level (survival stage). We use a Markov chain model to investigate the family level interactions (tactical decisions) and a life-history model to investigate the population level competition and the resource allocation problem related to this stage (strategic decisions). Figure 2 shows the relation of the two models and our main question.

**Figure 2.**
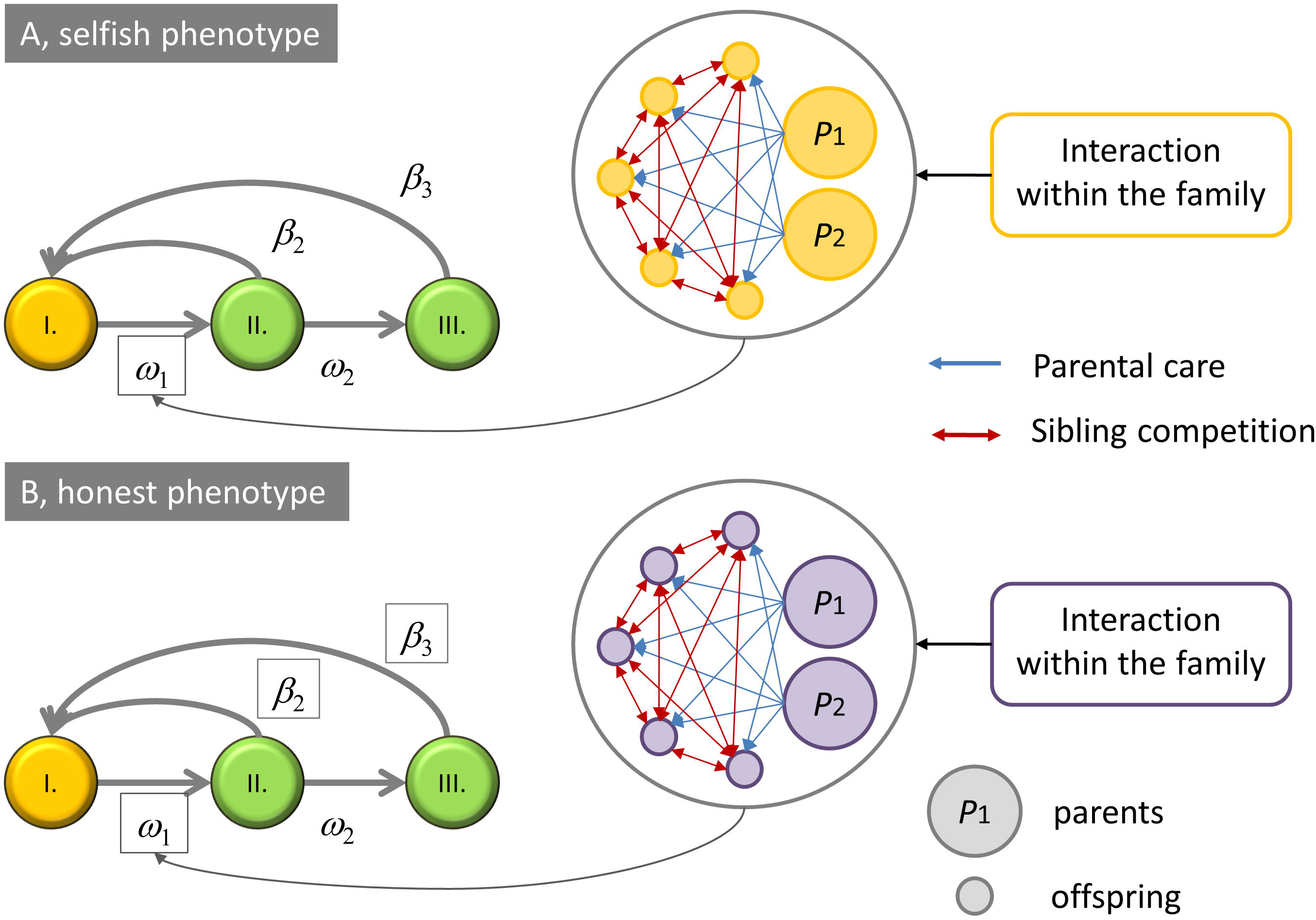
Relation of the tactical decisions made by the parents (Markov chain model) to the strategic decisions (life-history model) in the two-stage model. Temporal interactions within the family determine the survival rates of offspring (*ω_1_*), which is plugged into the life-history model (where *β* denotes fecundity).

**Figure 3.**
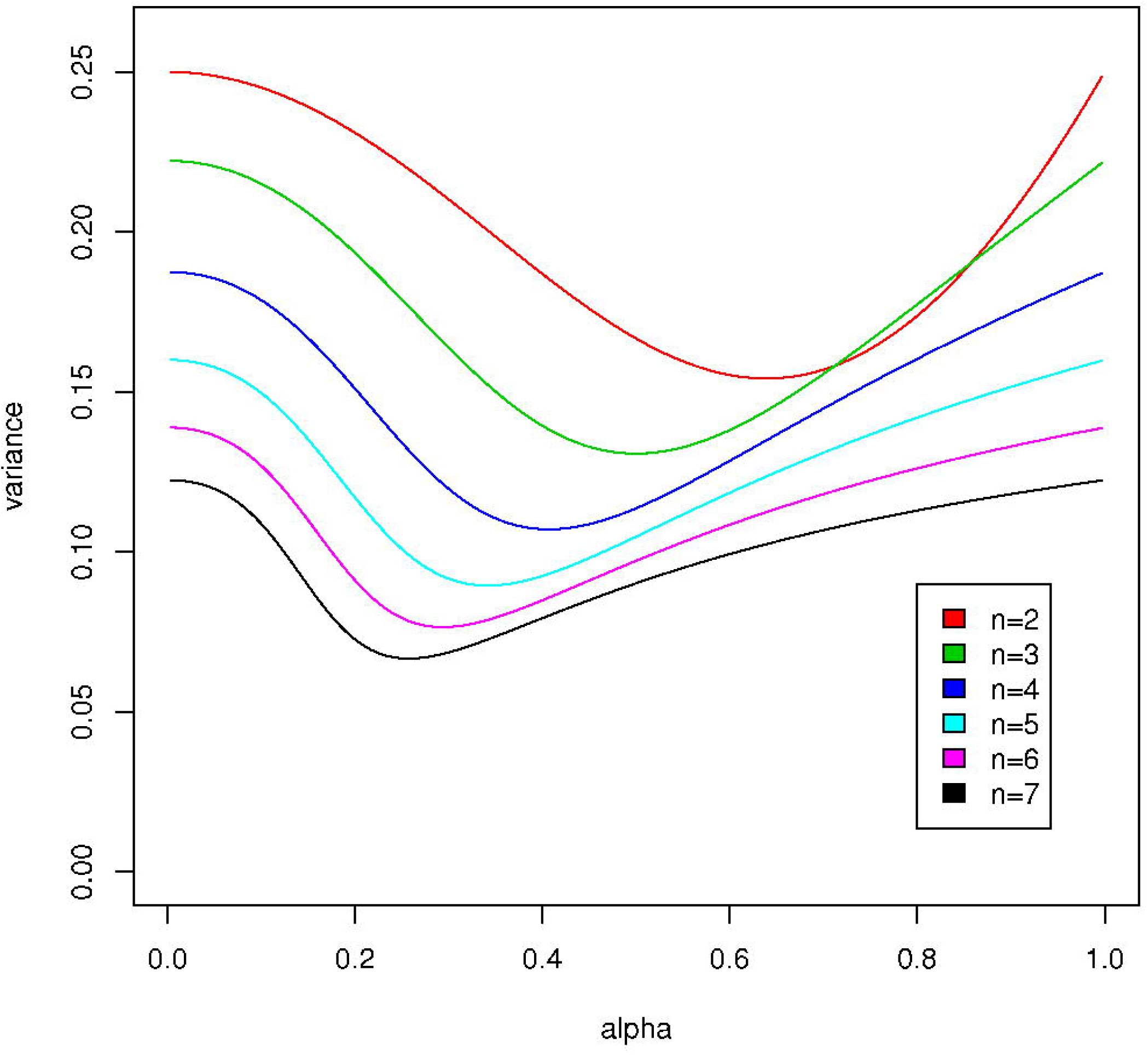
The asymptotic variance σ^2^(*α*), as a function of *α*, for offspring numbers *n* = 2 (top) to *n* = 7 (bottom).

We present a null model, which is the simplest model possible. We investigate whether honest begging could provide any advantage in such scenario. If such null model can show that honest begging can be advantageous under some conditions, then this makes other more complicated models unnecessary under the same conditions. Here we show that honest begging decreases the variance of food acquired by siblings, which leads to higher survival when the clutch size is around the optimum.

## 2. Results

First, we discuss the results of the dynamic temporal model (tactical level).

**Result 1**. The accumulated amount of the focal nestling’s food (*Y_T_*) can be approximated with a normal distribution.

**Result 2**. The variance (*Tσ*^2^(*α*)) of this normal distribution depends on two parameters: the number of nestlings (*n*) and the digesting parameter (*α*).

**Result 3**. The variance of the food amount acquired by the focal nestling in an honest family is always smaller than in the selfish family.

**Result 4**. If *m* > *m_f_*, then the survival probability of the focal nestling (*q*) is a monotone decreasing function of *σ*^2^(*α*), which means that if food is abundant enough, then the honest family will raise more surviving nestlings on average than the selfish family. On the other hand, if *m* < *m_f_*, then *q* is a monotone increasing function of *σ*^2^(*α*), which means that if food is scarce, then the selfish family will have the largest number of surviving offspring on average.

**Example 1**. Take the survival probability function *f*(*x*) with *m_f_* =280 and *σ_f_*=5. Figure 4 shows the mean number *E*(*V*) of surviving nestlings for different nest sizes *n*, as a function of the parameter *α* for fix *T* = 1500. Notice that when the number of nestlings is less than five, then food is abundant, and practically all nestlings are sure to survive. When the number of nestlings is more than five, then food is scarce, and practically all nestlings will die. The most interesting case is when there are 5 nestlings, all of which will get 300 units of food on average. In this case the expected number of survivors is between 4.45 and 4.72, depending on *α*.

**Figure 4.**
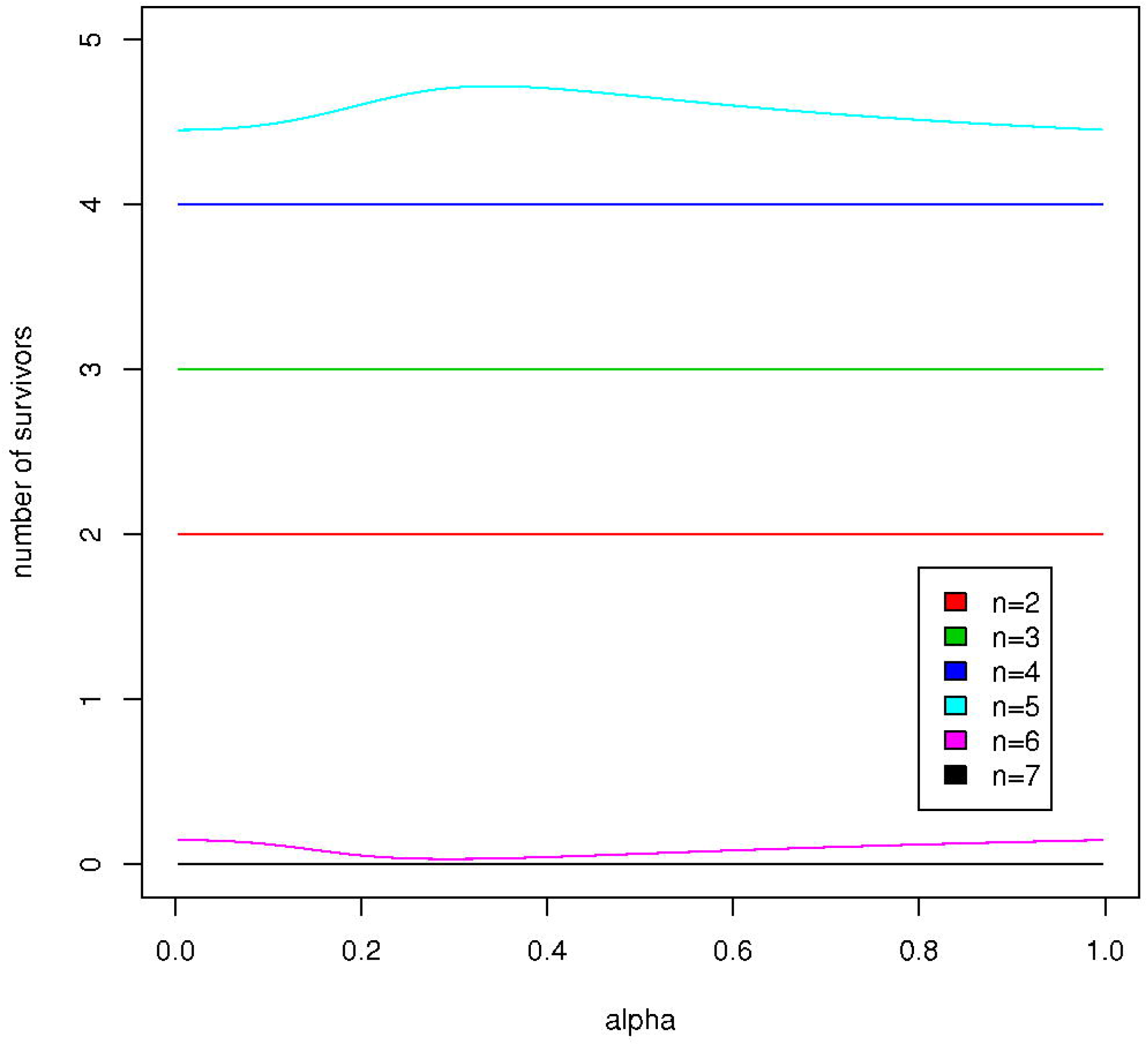
The mean number *E*(*V*) of surviving nestlings for different nest sizes *n*, as a function of the parameter *α*.

Next, we discuss the results of the resource allocation problem (strategic level). We investigate two situations: (i) a fixed amount *T* of food is distributed between nestlings; (ii) the total food *M* is fixed, and the mother distributes only the 0 < *s* < 1 fraction of it between nestlings (i.e. the total food distributed between nestlings is *T=sM*) and eats the remaining amount of *Θ* = (1 − *s*)*M*.

**Result 5**. Fixed *T*. The growth of population can be characterized by the leading eigenvalue *λ* of matrix *L*. This growth rate is a strictly increasing function of fecundities *β*(*α*) (see Remark 1), which in turn depend on the survival probability of the focal nestling *β*(*α*) *= nq*(*α*).

**Example 2**. It is seen that the characteristics of the curves are similar to the mean number *E*(*V*) of surviving nestling as a function of *α* (Fig. 5), thus the introduction of simple age structure does not alter qualitative outcome (cf. Fig. 4).

**Figure 5.**
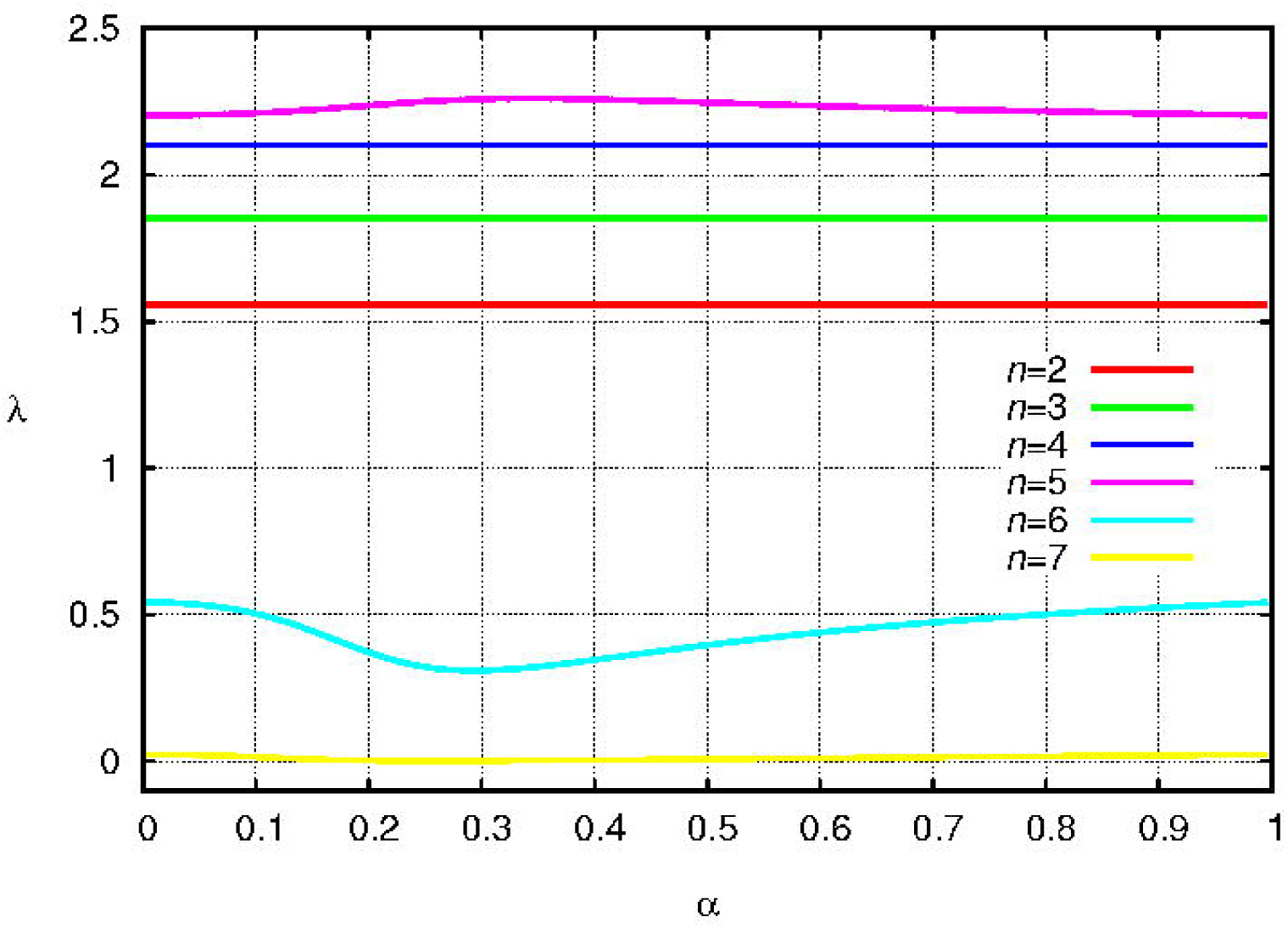
The leading eigenvalue of Leslie matrix *L* as a function of parameters *α* for different nest sizes *n*, with *T* = 1500.

**Result 6**. *T* depends on *s* (*T=sM*). In this case, the survival probability is a function of *s*, while the fecundity is a function of both *s* and *α β*(*α*, *s*). The growth rate of the population is a strictly increasing function of *β*(*α*, *s*).

**Example 3**. The leading eigenvalues as the functions of the two parameters *s* and *α* are shown in Fig. 6, upper panels (for *M* = 1800). The characteristics (near to the maximum in *s*) as a function of *α* are similar to the results of the simple model: in case of *n*=5 a maximum, if *n*=6 a minimum appear. Fig. 6 bottom panel shows the intersection curves of these graphs at *s*=0.82 (*n*=5) and at *s*=0.83 (*n*=6) (these *s* values correspond to the maximum of *λ* as a function of *s*).

**Figure 6.**
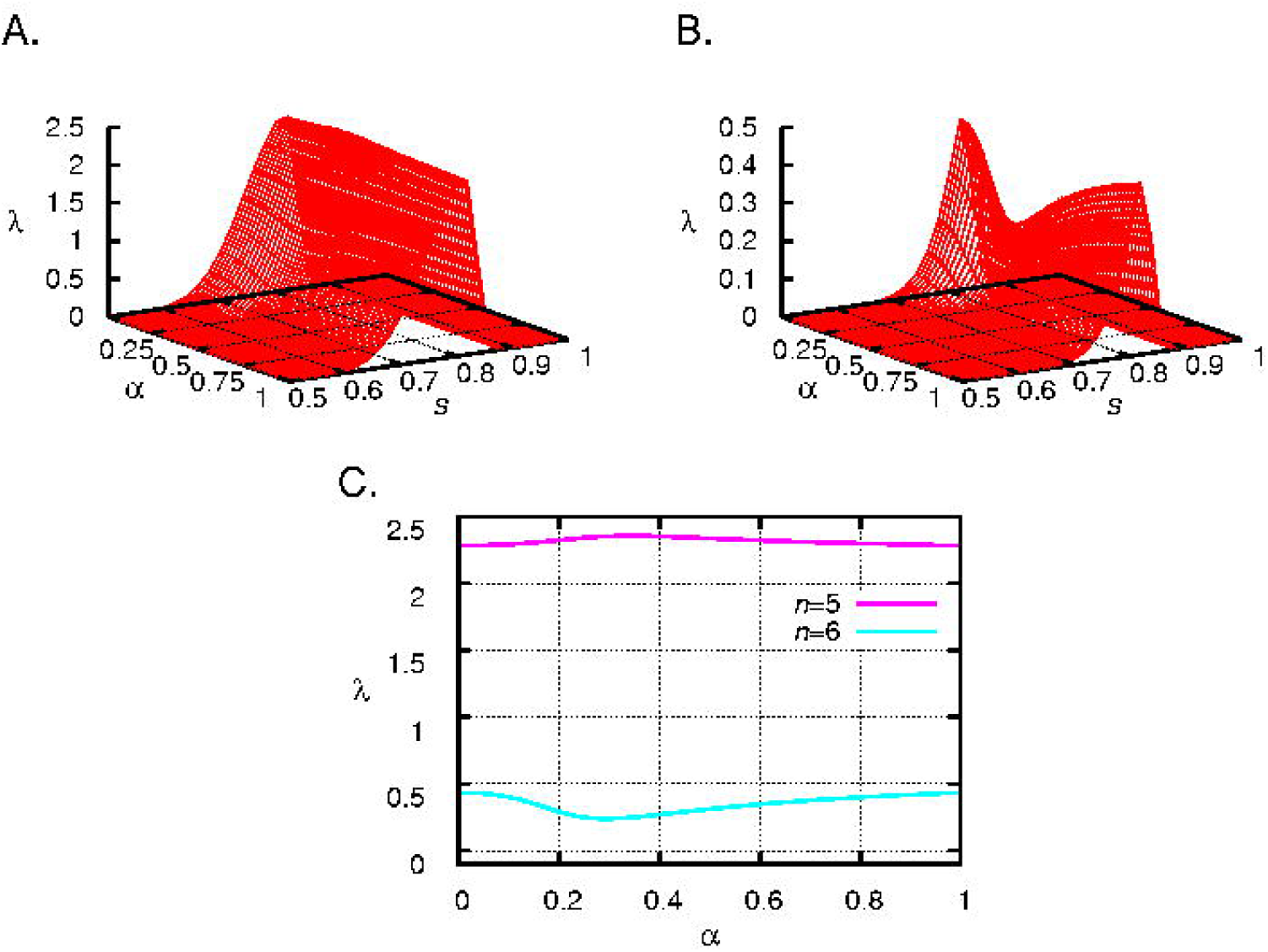
The leading eigenvalue as the function of *α* and *s* with different number of offspring, *n*=5 (top left), *n*=6 (top right). The leading eigenvalue as the function of *α* at *s*=0.815 with *n*=5 (bottom graph, red curve), at *s*=0.83 with *n*=6 (bottom graph, green curve). *M* = 1800. Furthermore, in the case of 5 offspring, for s<0.78 the selfish family, for s>0.78 the honest family performs better. Analogously, if *n*=6, for s<0.93 the selfish family, in the opposite case the honest family is better.

**Conclusion**. The introduction of age-structure in a biologically plausible way does not change the qualitative outcome of the dynamical model model.

Interestingly, in case of *n*=5 the maximum of the mean number of surviving nestlings (Fig. 4) and the maximum of the leading eigenvalues in both structured population models (Figs. 5 and 6) occurs at the same *α* (*α* ≈ 0.342). Similarly, if *n*=6 the position of minimum is the same for all three cases (*α* ≈ 0.293). This is again a consequence of Remark 1.

Note that, while the change in lambda as a function of alpha is relatively small (cf. Fig 6, especially in case of *n*=5), but this value characterized the growth rate of an exponential process. In this case small change in the growth rate results in large difference in population size, even if the time span is relatively short.

## 3. Models and Methods

### 1. A tactical model of offspring begging

Here we investigate the tactical decision making of the parents in the temporal dimension with the help of a Markov chain model of offspring begging. We model the begging and feeding behaviour of offspring and parents within a nest until fledging. At the end of the interaction we calculate the survival probability of the offspring based on the food accumulated during the period of parental care. We make three simplifying assumptions: (i) we assume that there is no quality difference between the offspring, (ii) we also assume that offspring has unlimited capacity (cannot be saturated by food) and (iii) they cannot die of hunger during the period of parental care. These assumptions are clearly unrealistic; however, our goal is to present the simplest analytically tractable model of offspring begging.

Each nest has *n* offspring. Parents bring a food item to the nest at every time step. Each food item has the same value, say 1. The reproductive stage consists of *T* time steps (*T* is supposed to be large enough), it follows that the parents bring *T* units of food to the nest. We further assume that parents keep some food to themselves (*M* – *T*) out of the total amount of food they gathered (*M*).

We consider the following stochastic model: each nestling can behave in two different ways: it opens its beak either wide (i.e., it begs for food) or narrow (i.e., it does not beg for food). The parent chooses a begging nestling at random, and puts the food in its beak. If none of the nestlings are begging, the parent chooses one of the *n* nestlings at random, and puts the food in its beak.

We suppose that after receiving food, a nestling will get hungry after a geometrically distributed amount of time, with parameter *α*. We will say that a nestling is digesting, if it is not hungry. If a nestling is digesting at time *t*, the probability that it is still digesting at time *t* + 1 is (1 − *α*).

We consider two phenotypes of families. In the selfish (or dishonest) family, all nestlings are begging for food all the time. In contrast, in the honest family, a nestling will beg only if it is hungry. It is worth noting that the selfish family is a special case of the honest family with *α* = 1.

Let us fix a survival function *f*, where *f*(*x*) denotes the probability that a nestling who received a total amount *x* of food up to time *T* will survive the reproductive stage. Our main goal is to find the expected value of surviving nestlings in each of the bird families.

To this end, we concentrate on a focal nestling in the nest, and calculate its survival probability q. This probability will depend on the parameters *n, T, α* and the function *f*. Since all nestlings behave in exactly the same way, it is immediate that the expected value of the focal nestling’s food *Y_T_* accumulated up to time *T* is *m* := E(*Y_T_*) = *T/n*. Note that for the selfish family (*α* = 1), *Y_T_* follows a binomial distribution of order *T* and parameter 1/*n*, thus it can be approximated with a normal distribution with expectation *m* and variance *Tσ*^2^(1), where 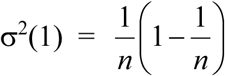. We show in Appendix 1 that this result holds more generally: for any 0 < *α* <1, *Y_T_* is approximately normally distributed with expectation *m* and variance *Tσ*^2^(*α*), and we give the precise analytic form of the asymptotic variance *σ*^2^(*α*). We can see that for any nestling number *n* and 0 < *α* < 1, the inequality 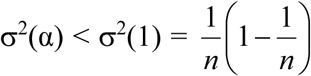 holds. Thus, we can conclude that in the honest family, the variance of the food amount acquired by the focal nestling is always smaller than in the selfish family.

Let us turn to the survival of the young birds. We have *n* nestlings, whose accumulated food amounts are *Y_T_*,_1_, *Y_T_*,_1_,…, *Y_T_*,*_n._* Denote by *V_i_* the indicator of the event that the *i*^th^ nestling survives. Then the number of surviving young birds is 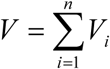 with expected value

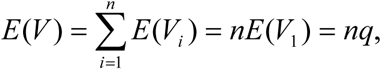
where *q* is the survival probability of the, say, focal nestling. If the survival function is *f*(*x*), and we use the normal approximation *Y_T_* ~ N(*m*, *Tσ^2^*(α)) for the accumulated food amount of the focal nestling, then

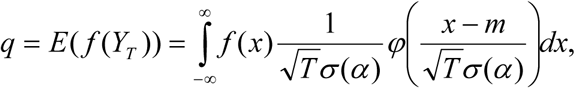
where φ(*x*) denotes the standard normal density. As an illustrating example, let 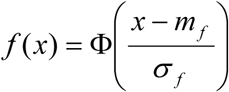, where Φ(*x*) denotes the standard normal distribution function. Here *m_f_* is the food amount which guarantees 50% survival probability: *f*(*m_f_*) = 0.5, and the *σ_f_* parameter regulates the „steepness” of *f* around *m_f_*. Let *U, W* be two independent standard normal variables, and 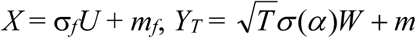. Then

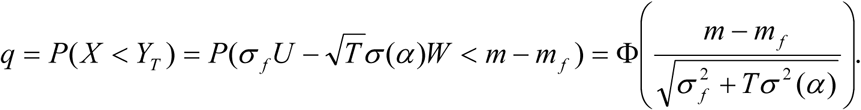

### 2. Kin demographic selection model

Here we investigate the strategic food allocation problem of the parents with a life-history model. We build on the general kin demographic selection model, introduced by Garay et al. [18]. The model is a life-history model with two stages during a year: reproduction and survival. We use the following assumptions: (i) different phenotypes do not interact during the reproductive stage: there are interactions only within the kin (here between offspring), (ii) different phenotypes have different hereditary life history parameters (juvenile survival rate, fecundity); and (iii) the survival rate depends on total density and it is the same for all individuals, independently of their phenotype and age; and finally (iv) the interactions within the kin during reproduction and the density dependent survival process during the survival stage are independent. Following these assumptions our model can be described as a combination of the following two sub-models.

The first sub-model describes what happens during the reproduction season, when the non-interacting females produce offspring, and the interactions (determining the demographic parameters of the different phenotypes) take place only within the family, i.e. between the offspring of the same given single female and between parents and their offspring. Here, the main point is that the phenotype-dependent demographic parameters determine the next age-classified state vector of the phenotype, so the stage of each phenotype is a demography vector, and each phenotype can be described by a Leslie matrix. Figure 7 shows an example.

**Figure 7.**
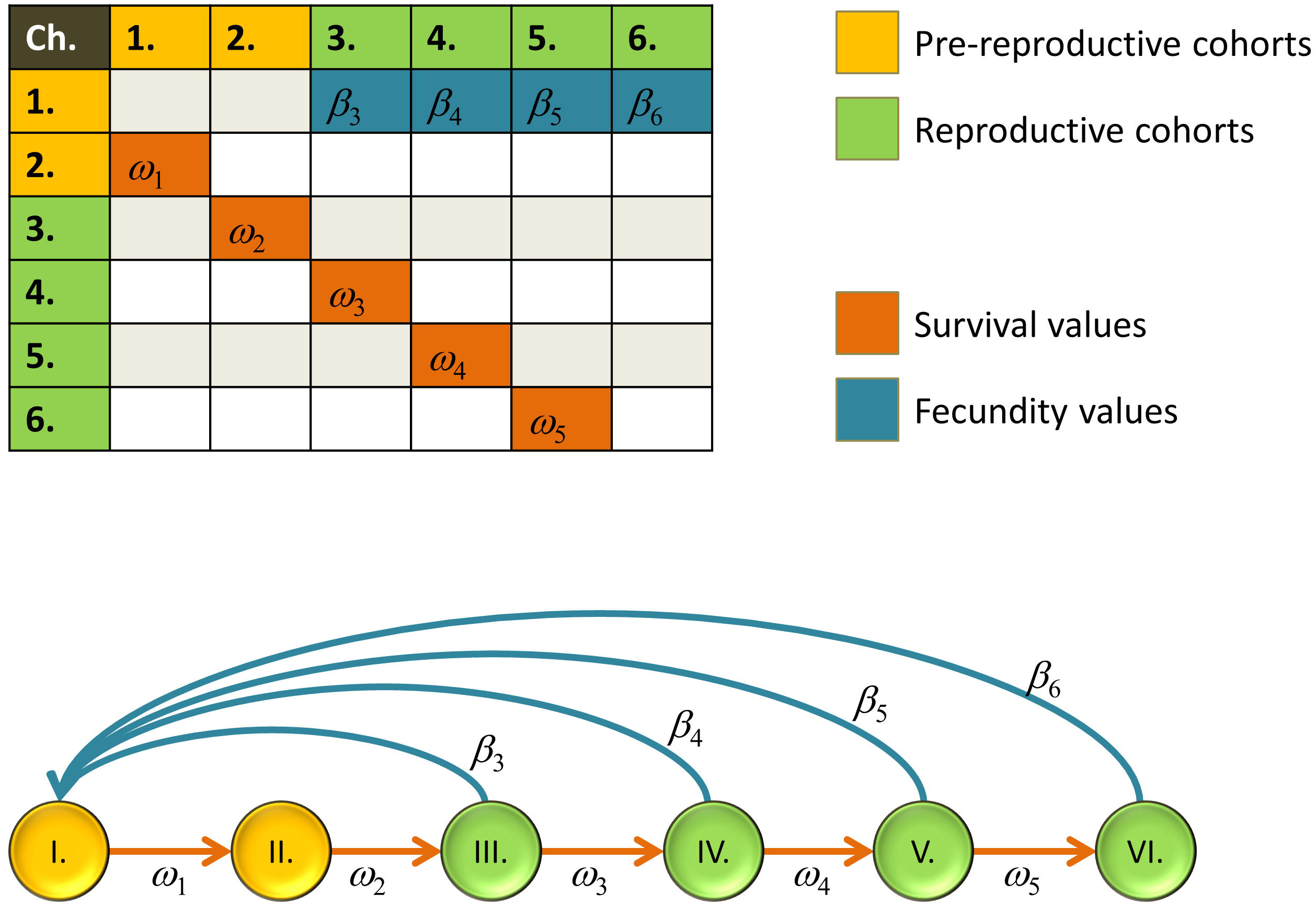
Life-history diagram and the corresponding Leslie matrix. Fecundity values and survival rates denoted by *β_i_* and *ω_i_* respectively.

Formally, consider a species with 3-year long life span, and each phenotype is characterized by a Leslie matrix of the form

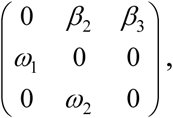
where either the fecundities *β_i_* or the survival rates *ω_i_*, or all of them depend on the behaviour phenotype, so in this case 4 parameters characterize a phenotype. These parameters are determined by what happens between family members during the reproductive season.

In the second sub-model, by a random survival process, the total population size is reduced to a fixed carrying capacity. Here, the main point is that the density-dependent survival process during autumn and winter, has a uniform effect on the demographic parameters of all individuals of all phenotypes. Formally, let us consider phenotypes *X* and *Y* having the respective population vectors *x*(*t*), *y*(*t*), developing according to Leslie models with respective matrices *L*_1_ and *L*_2_, the corresponding total densities are 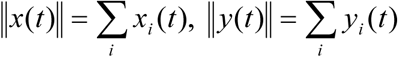, and phenotypic long-term growth rates are λ_1_ and *λ*_2_. The uniform survival by selection means that at given time *t*, each individual survival probability is proportional to

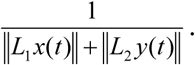

We have shown in our previous model (Garay et al., 2016) that if *λ*_1_*, λ*_2_ *>* 1 and *λ*_1_ *> λ*_2_, then

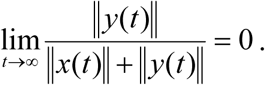

We have found that, in the framework kin demographic selection model, the fitness of the phenotype is the phenotypic long-term growth rate. This is the dominant positive eigenvalue of the Leslie matrix, which is a solution of the following characteristic polynomial:

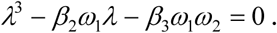

**Remark 1**. Clearly, the phenotypic long-term growth rate (λ) strictly increases in *β*_2_ and *β*_3_. Indeed, the positive solution of the characteristic polynomial *λ* satisfies

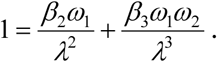

Hence 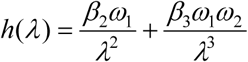 strictly increases in *β*_2_ and *β*_3_ thus λ also increases.

Now we will apply the above kin demographic selection model. The phenotype of an individual determines two behaviours in our model: its behaviour during the juvenile stage, and the behaviour during the adult stage. There are two potential trade-offs in the phenotypic Leslie matrix.

The first trade-off takes place in the parent-offspring interactions: parents have limited resources, they have to feed themselves and their offspring from a fix quantity of food. Accordingly, the parent’s phenotype determines both the survival rate and the fecundity parameter of the phenotypic Leslie matrix. The second one is in between the offspring: an honest nestling is begging food only when it is hungry, a selfish nestling is always begging food, thus the latter strategy decreases the survival rate of its sibs. Therefore, the phenotype of juveniles effects the fecundity parameters of the phenotypic Leslie matrix (see Section 3 and 4). We investigate two distinct phenotypes: honest and selfish. Since we consider an asexual population therefore either all juveniles are honest or all are selfish in any given nest. The reasons for this are the following: First, when a mutant appears, a mixed family is formed (there are both honest and selfish juveniles), but independently of mutant’s success in a mixed family, in the next generation all offspring of the mutant has the same mutant strategy in an asexual population. Secondly, concerning the long-term growth rate of phenotypes, the initial success of a single mutant is not so important.

To extend our investigation we introduced age-structure in populations. For sake of simplicity, we assume that neither the fecundity nor the survival do not depend on the age of the parent. Let us consider a 3 year-long life span described by the Leslie-matrix (*L*) with fecundities *β_i_ = β*, and survival rates *ω_i_ = ω*. (For sake of simplicity we assume same fecundities and same survival rates for the two age-classes.)

In the first scenario we assume that a fixed amount *T* of food is distributed between nestlings, and *M* is such that the survival probability of the parent is *ω* = 0.95. We assume that the fecundities are *β*(*α*) = *nq*(*α*), where *q*(*α*) is the survival probability of the focal nestling (see previous section). The growth of population can be characterized by the leading eigenvalue *λ* of matrix *L*. According to Remark 1, this growth rate is a strictly increasing function of *β*(*α*).

In the second scenario let us assume that the total food *M* is fixed, and the mother distributes only the 0 < *s* < 1 fraction of it between nestlings (the total food distributed between nestlings is *T*=*sM*) and eats the remaining amount of Θ = (1 − *s*)*M*. Let us assume that the survival probability function of a mother as a function of Θ has the following form: 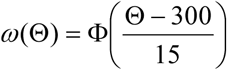.

In this case, the survival probability is a function of *s*, while the fecundity is a function of both *s* and *α*. According to Remark 1, for any fixed *s*, the leading eigenvalue is a strictly increasing function of *β*(*α, s*).

## 4. Conclusion

We found that the honest offspring’s solicitation for food over performs the selfish phenotype in asexual populations. Our result connects to basic view of idea of bet-hedging, namely the fitness variance is minimized by natural selection (see e.g. [19]). The classical objective (task) of bad-hedging theory how should individuals optimize their fitness in an unpredictable environment [20,21]. Different types of bet-hedging are listed in the literature like “conservative” [19,22] or “diversified” [23]. Individuals lower their expected fitness in both case in exchange for a lower variance either in their fitness (conservative) or in the survival of offspring (diversified). Contrary to these scenarios the source of randomness during parent-offspring interactions comes from the fact that the parent is not fully informed from the actual stage (either hungry or digesting) of its offspring in our case. Our result is different in another crucial way: the honest signal decreases the variance of fitness, and increases the mean of fitness, at a time. This is different from the traditional definition: “*Bet-hedging is defined as* a *strategy that reduces the temporal variance in fitness at the expense of* a *lowered arithmetic mean fitness*” [24], our result does not satisfy this strict definition of bet-hedging.

While the standard model of parent-offspring conflict corresponds to the sexually reproducing species [7] here we considered parent-offspring interaction in an asexual population. Thus, the direct theoretical comparison of these models is impossible. The main obstacle for the comparison is the following. Parent-offspring conflict there are two closely related individuals with different phenotypes, thus there are two objective functions: offspring’s and parents’ fitness in an asymmetric interaction. Contrary to that, in our model we have a two-dimensional life history strategy (what an individual does at its juvenile and the adult stage), and since the interaction happen within the family, thus our interaction is symmetric. The theoretical comparison of these models is an open problem.

We think that this kind of modelling approach is a necessary first step to resolve the debate about signals of hunger, need and quality and to gain a deeper understanding of food solicitation in the context of parent-offspring conflict.

## Acknowledgements.

This work was partially supported by the Hungarian National Research, Development and Innovation Office NKFIH [grant numbers K 108615 (to T.F.M.), K 108974 (to J.G. and SZ.SZ), K119347 (to A.SZ.) and GINOP 2.3.2–15–2016–00057 (to J.G., A.SZ)]. SZ.SZ. was supported by the European Research Council (ERC) under the European Union’s Horizon 2020 research and innovation programme (grant agreement No 648693).

## Appendix 1. Details of the Markov chain model

If 0 < *α* < 1, then we model the feeding process with a Markov chain *X_t_*, *t* = 1, 2, …, *T*, where *X_t_* denotes the state of the process just after the parent has given away the *t*^th^ unit of food (i.e., the state of the process at time *t*). We will have a total of *d* = 3*n* − 2 different states, each state will be a pair (*i, k*), where *i* denotes the state of the focal nestling: it can be any of the three possibilities *H* (hungry), *D* (digesting previously received food) or *R* (it just received food). The second coordinate *k* records the number of digesting nestlings (either digesting food received earlier, or food received at this moment) among the remaining *n* − 1 ones. There are *n* − 1 states (*H, k*), *k* = 1, …, *n* − 1, also *n* − 1 states (*D, k*), *k* = 1, …, *n* − 1, and *n* states (*R, k*), *k* = 0, 1, …, *n* − 1. We denote the state space by *S*. From the previous description of the process, it is immediate to write down the transition matrix *P* of the Markov chain *X_t_*. The elements of *P* are the transition probabilities *P*(*u, v*) = P(*X_t +_* _1_ *= v | X_t_ = u*), where *u, v* are any two states. Let us denote the binomial probabilities by 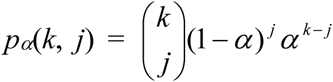 (this is the probability that out of *k* digesting nestlings, exactly *j* are still digesting at the next time step) and let 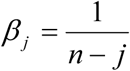. Then the (*H, k*)^th^ row of the transition matrix is:

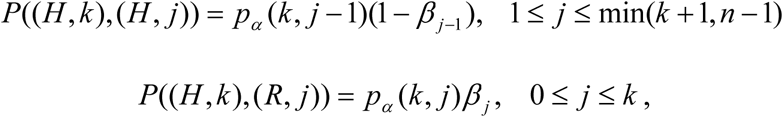
and all other elements are zero. The (*R, k*)^th^ row of the transition matrix for *k* < *n* − 1 is:

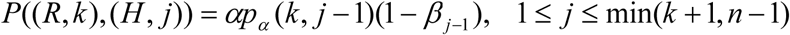

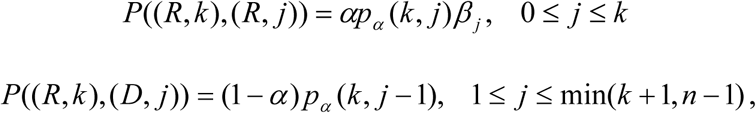
and all other elements are zero. The (*R, k*)^th^ row of the transition matrix for *k* = *n* − 1 is of the same form, except that

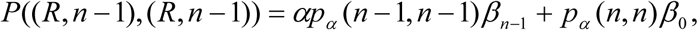
and

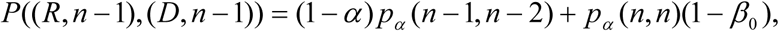
where the additional terms cover the cases when none of the nestlings are hungry. Finally, the (*D, k*)^th^ row of the transition matrix is exactly the same as the (*R, k*)^th^ row.

It is clear that our Markov chain is irreducible, and thus it is positive recurrent admitting a unique stationary distribution *π* = (*π_s_* : *s* ∈ *S*). Let *R\** = {(*R, k*): *k* = 0, 1, …, *n* − 1} be the subset of the states in which the focal nestling receives the food. Then the focal nestling’s accumulated amount of food *Y_T_* is just the number of visits of the chain in *R^*^* up to time *T*. By introducing the reward function on the states

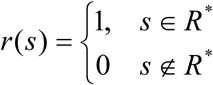
we can write 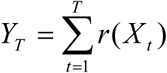. It is a standard result that the central limit theorem holds for *Y_T_*. To formulate the precise statement, let Π be the *d* × *d* matrix whose (*u, v*)^th^ entry is *π_v_*, and Π_dg_ the *d* × *d* diagonal matrix whose (*u, u*)^th^ entry is *π_u_*, and all other entries are 0. The *d* × *d* identity matrix is denoted by *I*. Finally, let *Z* = (*I* − *P* + Π)^−1^ be the so-called fundamental matrix of *P*.

### Theorem 1.

(Central limit theorem for Markov chains, e.g. Kemeny and Snell, 1969) Let *X_t_* be an irreducible Markov chain with finite state space *S*, and *r* any reward function on the states. Let *π* be the unique stationary distribution of the chain, and define

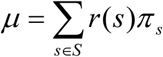
the mean of the reward function under stationarity, and

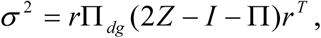
where *r* = (*r*(*s*): *s* ∈ *S*) is the row vector of the rewards. Then for any initial distribution of *X*_1_,

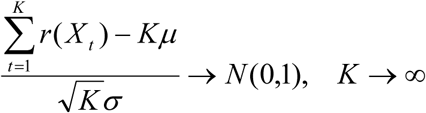
in distribution, where *N*(0, 1) stands for the standard normal distribution.

Let *A* be the *d × d* matrix with all its entries equal to 1, and **1** = (1, …, 1). Since *π* = **1**(*I* − *P + A*)^−1^, both *μ* and *σ* are straightforward to calculate from *P* by using matrix arithmetics.

In our case, as noted earlier,

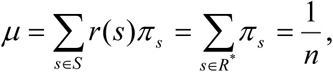
since in the stationary scenario, each nestling acquires the food with the same probability.

## References

1. Caro SM, Griffin AS, Hinde CA, West SA. Unpredictable environments lead to the evolution of parental neglect in birds. Nature communications. 2016;7:10985.

2. Caro SM, West SA, Griffin AS. Sibling conflict and dishonest signaling in birds. Proceedings of the National Academy of Sciences. 2016; 113(48): 13803–8.

3. Dugas MB. Baby birds do not always tell the truth. Proceedings of the National Academy of Sciences. 2016 Nov 21: 201616640.

4. Bebbington K, Kingma SA. No evidence that kin selection increases the honesty of begging signals in birds. Evolution Letters. 2017;1(3):132–7.

5. Mas F, Kölliker M. Maternal care and offspring begging in social insects: chemical signalling, hormonal regulation and evolution. Animal Behaviour. 2008;76(4):1121–31.

6. Kilner R, Johnstone RA. Begging the question: are offspring solicitation behaviours signals of need? Trends in Ecology & Evolution. 1997; 12(1):11–5.

7. Trivers RL. Parent-offspring conflict. Integrative and Comparative Biology. 1974;14(1):249–64.

8. Grafen A. Biological Signals as Handicaps. Journal of Theoretical Biology. 1990;144(4):517–46. doi: 10.1016/s0022-5193(05)80088-8.

9. Godfray HCJ. Signalling of need by offspring to their parents. Nature. 1991;352(6333):328–30.

10. Johnstone RA, Godfray HCJ. Models of begging as a signal of need. The Evolution of Begging. 2002: 1–20.

11. Nöldeke G, Samuelson L. How costly is the honest signaling of need? Journal of Theoretical Biology. 1999;197(4):527–39.

12. Mock DW, Dugas MB, Strickler SA. Honest begging: expanding from signal of need. Behavioral Ecology. 2011;22(5):909–17.

13. Hinde CA, Godfray HC. Quality, need, or hunger; begging the question. Behavioral Ecology. 2011 Jul 16;22(6):1147–8.

14. Johnstone RA, Kilner RM. New labels for old whines. Behavioral Ecology. 2011 Sep 1;22(5):918–9.

15. Wright J. Honest begging: signals of need, quality, and/or hunger? Behavioral Ecology. 2011 Sep 1;22(5):920–1.

16. Kölliker M. On the meaning of hunger and behavioral control in the evolution of honest begging. Behavioral ecology. 2011 Sep 1;22(5):919–20.

17. Bossan B, Hammerstein P, Koehncke A. We were all young once: an intragenomic perspective on parent-offspring conflict. Proc R Soc B; 2013: The Royal Society.

18. Garay J, Varga Z, Gamez M, Cabello T. Sib cannibalism can be adaptive for kin. Ecological Modelling. 2016;334:51–9. doi: 10.1016/j.ecolmodel.2016.05.001.

19. Olofsson H, Ripa J, Jonzén N. Bet-hedging as an evolutionary game: the trade-off between egg size and number. Proceedings of the Royal Society of London B: Biological Sciences. 2009;276(1669):2963–9.

20. Philippi T, Seger J. Hedging one’s evolutionary bets, revisited. Trends in Ecology & Evolution. 1989;4(2):41–4.

21. Starrfelt J, Kokko H. Bet-hedging—a triple trade-off between means, variances and correlations. Biological Reviews. 2012;87(3):742–55.

22. Garay J, Móri TF. Monogamy has a fixation advantage based on fitness variance in an ideal promiscuity group. Bulletin of mathematical biology. 2012;74(11):2676–91.

23. Grantham ME, Antonio CJ, O’Neil BR, Zhan YX, Brisson JA. A case for a joint strategy of diversified bet hedging and plasticity in the pea aphid wing polyphenism. Biology letters. 2016;12(10):20160654.

24. Ripa J, Olofsson H, Jonzén N. What is bet-hedging, really? Proceedings of the Royal Society of London B: Biological Sciences. 2010;277(1685):1153–4.

